# Effects of Δ9-tetrahydrocannabinol (THC) vapor inhalation in Sprague-Dawley and Wistar rats

**DOI:** 10.1101/541003

**Authors:** Michael A. Taffe, K. M. Creehan, Sophia A. Vandewater, Tony M. Kerr, Maury Cole

**Author notes:** Address Correspondence to: Dr. Michael A. Taffe, Department of Neuroscience, SP30-2400; 10550 North Torrey Pines Road; The Scripps Research Institute, La Jolla, CA 92037; USA.

## Abstract

A novel inhalation system based on e-cigarette technology has been recently shown to produce hypothermic and anti-nociceptive effects of Δ^9^-tetrahydrocannabinol (THC) in rats. Indirect comparison of some prior investigations suggested differential impact of inhaled THC between Wistar (WI) and Sprague-Dawley (SD) rats, thus this study was conducted to directly compare the strains.

Groups (N=8 per strain) of age matched male SD and WI rats were prepared with radiotelemetry devices to measure temperature and then exposed to vapor from the propylene glycol (PG) vehicle or THC (25, 100, 200 mg/mL of PG) for 30 or 40 minutes. Additional studies evaluated plasma THC levels and anti-nociceptive effects after THC inhalation as well as the thermoregulatory effect of intraperitoneal injection of THC (5-30 mg/kg).

Hypothermic effects of inhaled THC was more pronounced in SD rats however plasma levels of THC were identical across strains under either fixed inhalation conditions or injection of a mg/kg equivalent dose. Strain differences in hypothermia were even more pronounced after i.p. injection of THC with SD rats exhibiting dose-dependent temperature reduction after 5 or 10 mg/kg, i.p. and the WI rats only exhibiting significant hypothermia after 20 mg/kg, i.p. The anti-nociceptive effects of inhaled THC (100, 200 mg/mL) did not differ significantly across the strains. These studies confirm an insensitivity of WI rats, compared with SD rats, to the hypothermia induced by THC following inhalation conditions that produced identical plasma THC and anti-nociception. Thus strain differences were not due to differential THC delivery via vapor inhalation.

## 1. Introduction

Humans increasingly use e-cigarette type devices filled with cannabis extracts to administer an active dose of Δ^9^-tetrahydrocannabinol (THC) and other constituents (Mammen et al. 2016; Morean et al. 2015; Morean et al. 2017). This has spurred development of pre-clinical models which are capable of a similar route of drug administration in laboratory rodents. Recent studies showed that intrapulmonary delivery of THC using an e-cigarette based system results in a robust, dose-dependent hypothermia and anti-nociceptive effect in male and female rats (Javadi-Paydar et al. 2018a; Nguyen et al. 2016) and a similar system results in hypothermia in mice following inhalation exposure to synthetic cannabinoid agonists (Lefever et al. 2017). Recent discussion of replication and reliability across many scientific disciplines identifies the generalization of effects beyond narrowly constrained experimental protocols as a key issue. It is therefore of significant interest to determine where, for example, rat strain does or does not affect experimental outcomes. Such differences may be qualitative (present in one strain, absent in another) or quantitative (differing in maximum extent of a response or in a dose-response relationship). While our prior studies of cannabinoid inhalation have shown efficacy in both Wistar and Sprague-Dawley rats, this has not yet been directly compared across the strains in a well controlled manner. The major goal of this study was to assess any strain differences using age- and treatment-matched male rats.

Previous studies have reported strain-related differences in the effects of THC in rats. For example, adolescent THC exposure differentially affects adult measures of learning with Wistar rats being less sensitive than Long-Evans rats (Keeley et al. 2015). Repeated adolescent THC injection resulted in different effects of heroin conditioned place preference in Fischer 344 versus Lewis rats (Cadoni et al. 2015). Strain differences are not always found, for example, there was no difference between the Fischer and Lewis rat strains in conditioned taste avoidance and the hypothermic effects of THC (Wakeford and Riley 2014).

A substantial and sustained decrease in body temperature, and a decrease in nociceptive sensitivity, are major indicators of cannabinoid-like activity in laboratory rodents (Wiley et al. 2014) and are therefore ideal *in vivo* readouts for determining efficacy. Prior studies from this laboratory found that vapor inhalation of THC using an e-cigarette based system decreases body temperature, with nadirs similar to the effects of 10-20 mg/kg THC, i.p. (Javadi-Paydar et al. 2018a; Nguyen et al. 2016) but with a duration of action that was much shorter (Taffe et al. 2015). The inhalation studies also found anti-nociceptive effects of inhaled THC that were comparable in magnitude to those produced by THC injection. Although we’ve previously shown efficacy of the THC inhalation procedure independently in Sprague-Dawley and Wistar strains, there appeared to be a slight difference in thermoregulatory sensitivity with Sprague-Dawley rats reaching a lower nadir body temperature (Javadi-Paydar et al. 2018a; Nguyen et al. 2016). Pilot studies for previously conducted i.p. injection studies (Taffe et al. 2015) found Wistar rats relatively insensitive to a given dose of THC but this was never followed up in any rigorous manner. Since our prior studies were not designed as direct strain-comparisons (i.e., with age-matched groups treated identically), additional investigation is needed. This study was therefore designed to determine if the body temperature and anti-nociceptive responses to the inhalation of THC via e-cigarette type technology differed across two common strains of laboratory rat.

## 2. Method

### 2.1 Subjects

Male Wistar (Charles River, New York; N=10) and Sprague-Dawley (Harlan, Livermore, CA; N=10) rats were housed in humidity and temperature-controlled (23±2 °C) vivaria on 12:12 hour light:dark cycles. Rats had *ad libitum* access to food and water in their home cages and all experiments were performed in the rats’ scotophase. Rats entered the laboratory at 9 weeks of age on a single arrival day. The majority of the studies were conducted on a fixed subset of N=8 per group with the remaining N=2 per group being used in a subset of the plasma studies, as described below. All procedures were conducted under protocols approved by the Institutional Animal Care and Use Committee of The Scripps Research Institute.

### 2.2 Radiotelemetry

Rats were implanted with sterile radiotelemetry transmitters (TA-F40; Data Sciences International, St Paul, MN) in the abdominal cavity as previously described (Taffe et al. 2015; Wright et al. 2012) at 10 weeks of age. For experiments, starting at 12 weeks of age, rats were evaluated in clean standard plastic homecages (thin layer of bedding) in a dark testing room, separate from the vivarium, during the (vivarium) dark cycle. Radiotelemetry transmissions were collected via telemetry receiver plates (Data Sciences International, St Paul, MN; RPC-1 or RMC-1) placed under the cages as described in prior investigations (Aarde et al. 2013; Miller et al. 2013). Test sessions for inhalation studies started with a 15 minute interval to ensure data collection, then a 15 minute interval for baseline temperature and activity values followed by initiation of vapor sessions in a separate chamber; animals were returned to the recording chamber after cessation of the inhalation interval. The 15 minute baseline was omitted for the studies with intraperitoneal injection of THC due to the delayed onset of hypothermia (Nguyen et al. 2016; Taffe et al. 2015). Pre-treatment drugs or vehicle for the antagonist studies were injected prior to THC administration as specified in the following experimental descriptions.

### 2.3 Drugs

Δ^9^-tetrahydrocannabinol (THC) was administered by vapor inhalation with doses described by the concentration in the propylene glycol (PG) vehicle and by the duration of inhalation. Our prior reports show that 30 minutes of inhalation of vapor produced from THC concentrations of 50-200 mg/mL produce plasma THC levels similar to those produced by 3-10 mg/kg THC, i.p. (Javadi-Paydar et al. 2018a; Nguyen et al. 2016). THC was also administered intraperitoneally in doses of 5, 10, 20 or 30 mg/kg. SR141716 (Rimonabant; SR) was administered intraperitoneally in a dose of 4 mg/kg. For injection, THC or SR were suspended in a vehicle of 95% ethanol, Cremophor EL and saline in a 1:1:8 ratio. The THC was provided by the U.S. National Institute on Drug Abuse; SR141716 was obtained from ApexBio (New Delhi, Delhi, India; Distributor: Fisher Scientific, Pittsburgh, PA, USA).

### 2.4 Inhalation Apparatus

Sealed exposure chambers were modified from the 259 mm × 234 mm × 209 mm Allentown, Inc (Allentown, NJ) rat cage to regulate airflow and the delivery of vaporized drug to rats as has been previously described (Nguyen et al, 2016a; Nguyen et al, 2016b). An e-vape controller (Model SSV-1; La Jolla Alcohol Research, Inc, La Jolla, CA, USA) was triggered to deliver the scheduled series of puffs from Protank 3 Atomizer (Kanger Tech; Shenzhen Kanger Technology Co.,LTD; Fuyong Town, Shenzhen, China) e-cigarette cartridges by MedPC IV software (Med Associates, St. Albans, VT USA). The chamber air was vacuum-controlled by a chamber exhaust valve (i.e., a “pull” system) to flow room ambient air through an intake valve at ~1 L per minute. This also functioned to ensure that vapor entered the chamber on each device triggering event. The vapor stream was integrated with the ambient air stream once triggered.

### 2.5 Plasma THC analysis

Blood samples were collected (~500 μl) via jugular needle insertion following anesthesia with an isoflurane/oxygen vapor mixture (isoflurane 5% induction). Plasma THC content was quantified using fast liquid chromatography/mass spectrometry (LC/MS) adapted from (Irimia et al. 2015; Lacroix and Saussereau 2012; Nguyen et al. 2017). 50 μl of plasma were mixed with 50 μl of deuterated internal standard (100 ng/ml CBD-d3 and THC-d3; Cerilliant), and cannabinoids were extracted into 300 uL acetonitrile and 600 μl of chloroform and then dried. Samples were reconstituted in 100 μl of an acetonitrile/methanol/water (2:1:1). Separation was performed on an Agilent LC1100 using an Eclipse XDB-C18 column (3.5um, 2.1mm × 100mm) using gradient elution with water and methanol, both with 0.2 % formic acid (300 μl/min; 73-90%). Cannabinoids were quantified using an Agilent MSD6140 single quadrodpole using electrospray ionization and selected ion monitoring [CBD (m/z=315.2), CBD-d3 (m/z=318.2), THC (m/z=315.2) and THC-d3 (m/z=318.2)]. Calibration curves were conducted daily for each assay at a concentration range of 0-200 ng/mL and observed correlation coefficients were 0.999.

### 2.6 Data Analysis

The telemeterized body temperature and activity rate (counts per minute) were collected on a 5-minute schedule in telemetry studies, but are expressed as 30 minute averages for analysis (baseline values are from 15 minute intervals). The time courses for data collection are expressed relative to the THC injection time or the initiation of vapor inhalation, and times in the figures refer to the end of the interval (e.g. “60 minutes” reflects the data collected from 35 to 60 minutes, inclusive). Any missing temperature values were interpolated from the values before and after the lost time point. Activity rate values were not interpolated because 5-minute to 5-minute values can change dramatically, thus there is no justification for interpolating. Data (telemeterized temperature and activity measures, tail withdrawal latency, plasma THC) were analyzed with Analysis of Variance (ANOVA) including repeated measures for the Drug treatment condition and the Time after vapor initiation or injection. Any significant main effects were followed with post-hoc analysis using Tukey (multi-level factors) or Sidak (two-level factors) correction. All analysis used Prism 7 for Windows (v. 7.03; GraphPad Software, Inc, San Diego CA).

### 2.7 Experiments

The timing of the experiments is schematized in **Figure 1** which also depicts the body weight of the groups across the study. Animals were exposed to THC no more frequently than weekly across the entire study.

**Figure 1:**
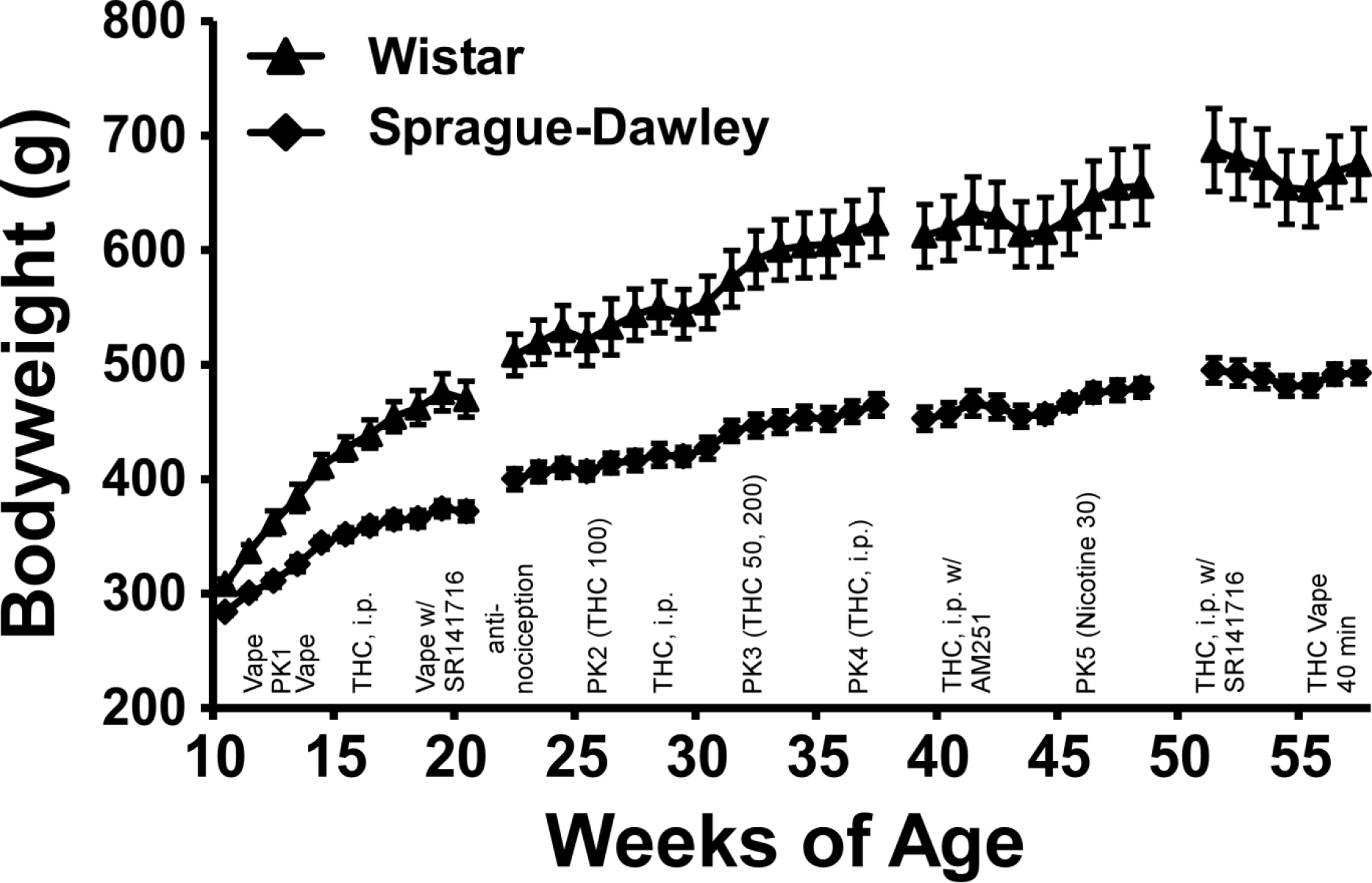
Mean (N=8 / group; SEM) body weight of the Wistar and Sprague-Dawley rats from 11 to 57 weeks of age. The approximate experimental timeline is also outlined, refer to Methods for full specification. PK indicates timing of blood draws for plasma assessment.

#### 2.7.1 Experiment 1 (Hypothermic response to Inhaled THC)

The inhalation sessions were 30 minutes in duration with four vapor puffs delivered every 5 minutes. The THC concentration was 0, 25 or 100 mg/mL with dosing order randomized within cohorts of 8 (N=4 per strain). The first cohort was evaluated before the first pharmacokinetic study (see below) and the second cohort was evaluated in weeks 14-15, with active doses being evaluated no more frequently than once per week.

#### 2.7.2 Experiment 2 (Plasma THC 1,2)

The rats were exposed to an inhalation session (THC 100 mg/mL for 30 minutes) after which a blood sample (~500 uL) was obtained via jugular venipuncture under anesthesia with an isoflurane/oxygen vapor mixture (isoflurane 5% induction) in week 13 (time 1) and in week 25 (time 2).

#### 2.7.3 Experiment 3 (Hypothermic response to THC, i.p.)

The rats were injected with THC (0.0, 5.0, 10.0 mg/kg, i.p.) in a counterbalanced order across weeks 16-18 with all animals tested concurrently. Due to the expected timecourse of effects (Taffe et al. 2015), there was no habituation interval beyond the initial 5-10 minutes to ensure telemetry sampling was operating properly. A followup study was conducted during weeks 27-28 in which rats were injected with a higher dose of THC (0.0, 20.0 mg/kg, i.p.) in a counter-balanced order within the groups.

#### 2.7.4 Experiment 4 (SR141716 prior to vapor inhalation)

A study was conducted in which rats were injected with the vehicle or SR141716 (4 mg/kg, i.p.) fifteen minutes prior to the start of 30 minute inhalation sessions of PG or THC (100 mg/mL) vapor in a counterbalanced order. Telemetry data were only available from 7 animals per group in this study due to experimental exigencies. Approximately half of the test days (counterbalanced order within each group) were completed prior to Experiment 5 and half after the second pharmacokinetic experiment.

#### 2.7.5 Experiment 5 (Anti-nociception)

The rats were exposed to inhalation sessions (PG, THC 100 mg/mL or THC 200 mg/mL for 30 minutes) and assessed on a tail-withdrawal assay (52 °C water bath) using methods previously described (Javadi-Paydar et al. 2018b). Tail withdrawal latency was assessed before the start of inhalation, after the session (35, 60 and 120 minutes after the start of inhalation).

#### 2.7.6 Experiment 6 (Plasma THC 3,4)

Blood samples were obtained in Weeks 33 and 34 following inhalation of THC (50, 200 mg/mL for 30 minutes in balanced order) and in Weeks 37 and 38 one hour after injection of THC (10, 20 mg/kg, i.p. in balanced order). (The groups also received vapor inhalation of nicotine (30 mg/mL) for 30 minutes in Week 46 for a plasma study not reported herein.) The additional 2 animals per group were included for these plasma experiments after participating in unrelated pilot experiments involving inhalation of oxycodone and heroin.

#### 2.7.7 Experiment 7 (SR141716 prior to THC i.p.)

The effect of pretreatment with vehicle versus the CB_1_ receptor antagonist/inverse agonist SR141716 (4 mg/kg, i.p.) on the body temperature response to vehicle or THC injected i.p. was assessed in Weeks 51-53. For this study the Wistar rats received a 30 mg/kg THC dose and the Sprague-Dawley rats received a 20 mg/kg dose. The reason for this difference was that a similar study conducted with the CB_1_ receptor antagonist AM251 (4 mg/kg, i.m.) in Weeks 41-42 found no hypothermia after 20 mg/kg THC i.p. in the Wistar rats (and no effect of AM251 on the response to 20 mg/kg THC, i.p. in the Sprague-Dawley group).

#### 2.7.8 Experiment 8 (THC vapor inhalation for 40 minutes)

The effect of inhalation of THC (100 mg/mL) for 40 minutes on body temperature was assessed at 55-56 weeks of age. The goal was to determine if the apparent insensitivity of the WI group (compared with the SD group and their own response to 20 mg/kg in Experiment 3) during the injection studies would also be reflected in THC inhalation at this age.

## 3. Results

The two strains differed significantly in bodyweight, with the WI group consistently heavier across the entire study (**Figure 1**). The statistical analysis confirmed significant effects of Age [F (42, 588) = 241.6; P<0.0001], Strain [F (1, 14) = 28.14; P<0.0005] and the interaction of Strain with Age [F (42, 588) = 18.35; P<0.0001] on bodyweight. The post-hoc test confirmed that the strains were significantly different from 17-56 weeks of age.

### 3.1 Experiment 1 (Hypothermic response to Inhaled THC)

The inhalation of THC (25 or 100 mg/mL) for 30 minutes reduced body temperature of each strain of rat (**Figure 2**). The statistical analysis using a combined between-groups factor for Strain and Drug Treatment confirmed significant effects of Time [F (7, 294) = 50.13; P<0.0001], Strain/Drug Treatment condition [F (5, 42) = 17.82; P<0.0001] and the interaction of factors [F (35, 294) = 10.67; P<0.0001] on the temperature. The Tukey post-hoc test confirmed that temperature of the strains differed from each other 60, 120 and 150 minutes after the start of THC inhalation (other post-hoc findings are depicted on Figure 2).

**Figure 2:**
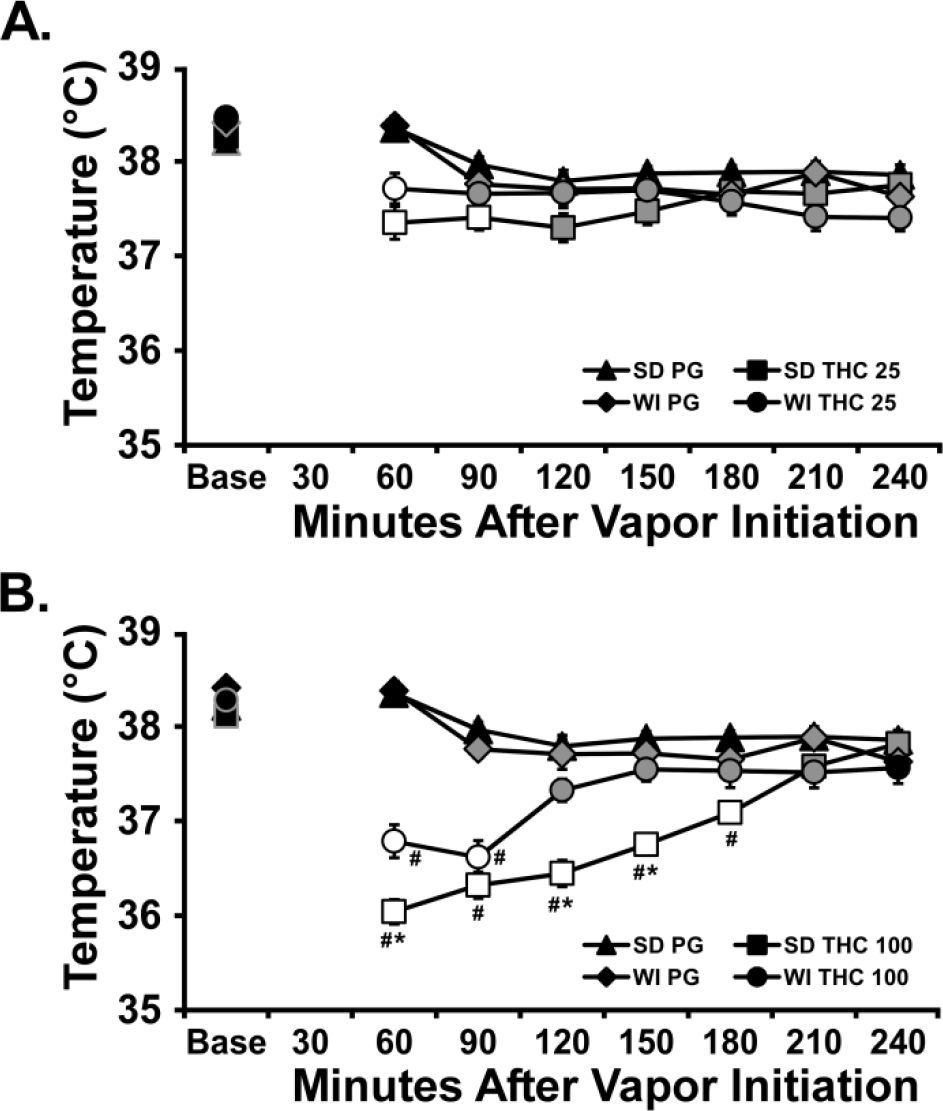
Mean (N=8 / group; SEM) body temperature following vapor inhalation of the PG vehicle or A) THC (25 mg/mL in the PG) or B) THC (100 mg/mL), for 30 minutes. The PG data are duplicated on upper and lower graphs for clarity. Base=baseline. Open symbols indicate a significant difference from both vehicle at a given time-point and the within-treatment baseline, while shaded symbols indicate a significant difference from the baseline only. A significant difference from the THC 25 mg/mL at a given time is indicated by # and between strains at a given time for the same dose by *.

A follow up three-way ANOVA confirmed significant effects of factors for Strain [F (1, 224) = 11.64; P<0.001], Time after the start of Initiation [F (7, 224) = 30.87; P<0.0001] and the Drug treatment condition [F (1, 224) = 294; P<0.0001], as well as all interactions of factors on the body temperature. There was a significant effect of Time [F (7, 294) = 80.54; P<0.0001], but not of Drug treatment condition or any interaction, on activity.

### 3.2 Experiment 2 (Plasma THC 1,2)

The inhalation of THC (100 mg/mL in the PG) for 30 minutes resulted in identical plasma THC between the two groups of rats (**Figure 3**) when assessed at 13 or 25 weeks of age, but a significantly higher plasma THC concentration was observed at the later time. A two-way ANOVA confirmed a significant effect of Week of sampling [F (1, 14) = 23.92; P<0.0005] but not of Strain or the interaction of factors. Examination of the individual values confirms the substantial overlap of the plasma THC levels reached in the two strains at either 13 or 25 weeks.

**Figure 3:**
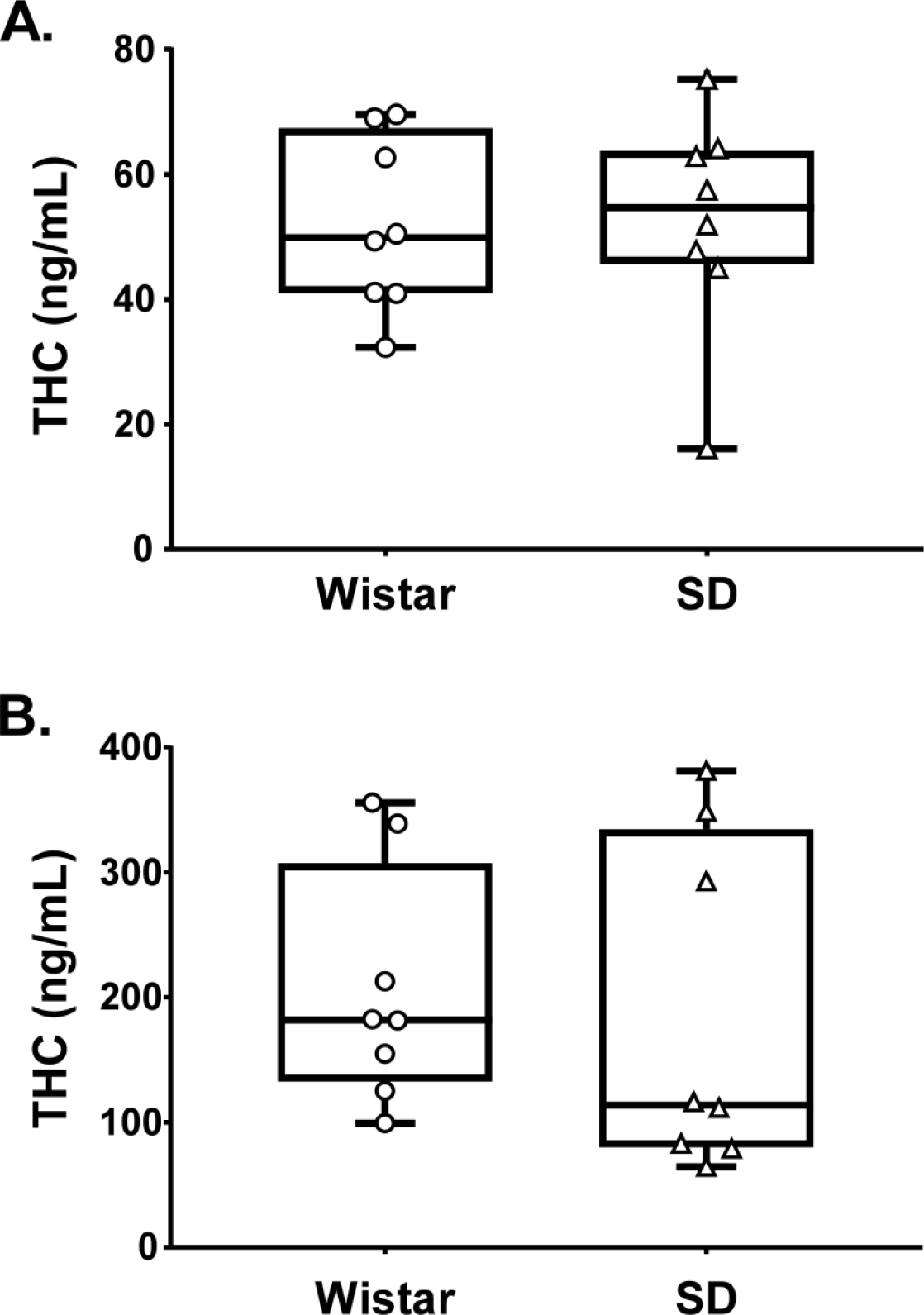
Plasma THC concentrations for Wistar and Sprague-Dawley rats (N=8 / group) following vapor inhalation of the THC (100 mg/mL) for 30 minutes at A) 13 and B) 25 weeks of age. Box plots depict median and interquartile range and the bars indicate the range. Individual subject values are also plotted.

### 3.3 Experiment 3 (Intraperitoneal Injection)

The groups were first injected with THC (0.0, 5, 10 mg/kg, i.p.) in a counter-balanced order and monitored for body temperature and activity levels. There was significant hypothermia in Sprague-Dawley but not Wistar rats (**Figure 4 A,B**), and the statistical analysis confirmed significant effects of Time [F (10, 420) = 33.56; P<0.0001], Drug treatment / Group [F (5, 42) = 7.152; P<0.0001] and the interaction of factors [F (50, 420) = 3.89; P<0.0001] on temperature after i.p. injection. Wistar and SD rat temperature differed significantly from each other after 10 mg/kg THC (150-300 minutes post-injection). The followup ANOVA within the SD group confirmed significant effects of Time [F (10, 70) = 22.84; P<0.0001], Drug treatment [F (2, 14) = 15.31; P<0.0005] and the interaction of factors [F (20, 140) = 6.87; P<0.0001] on temperature. The post hoc test confirmed that temperature was lower than the baseline in the SD rats after 5 mg/kg (120-300 minutes post-injection) or 10 mg/kg (60-300 minutes post-injection). Temperature was also significantly lower compared with the corresponding timepoint after vehicle in the SD rats after 5 mg/kg (120-300 minutes post-injection) or 10 mg/kg (60-300 minutes post-injection) and temperature differed significantly after the 5 mg/kg versus 10 mg/kg doses (60-270 minutes post-injection). The followup repeated measures ANOVA in the Wistar group confirmed a significant effect of Time [F (10, 70) = 19.31; P<0.0001], but not of Drug treatment or the interaction of factors on body temperature. The post hoc test confirmed that temperature was lower than the baseline in the WI rats after 10 mg/kg (60-210 minutes post-injection).

**Figure 4:**
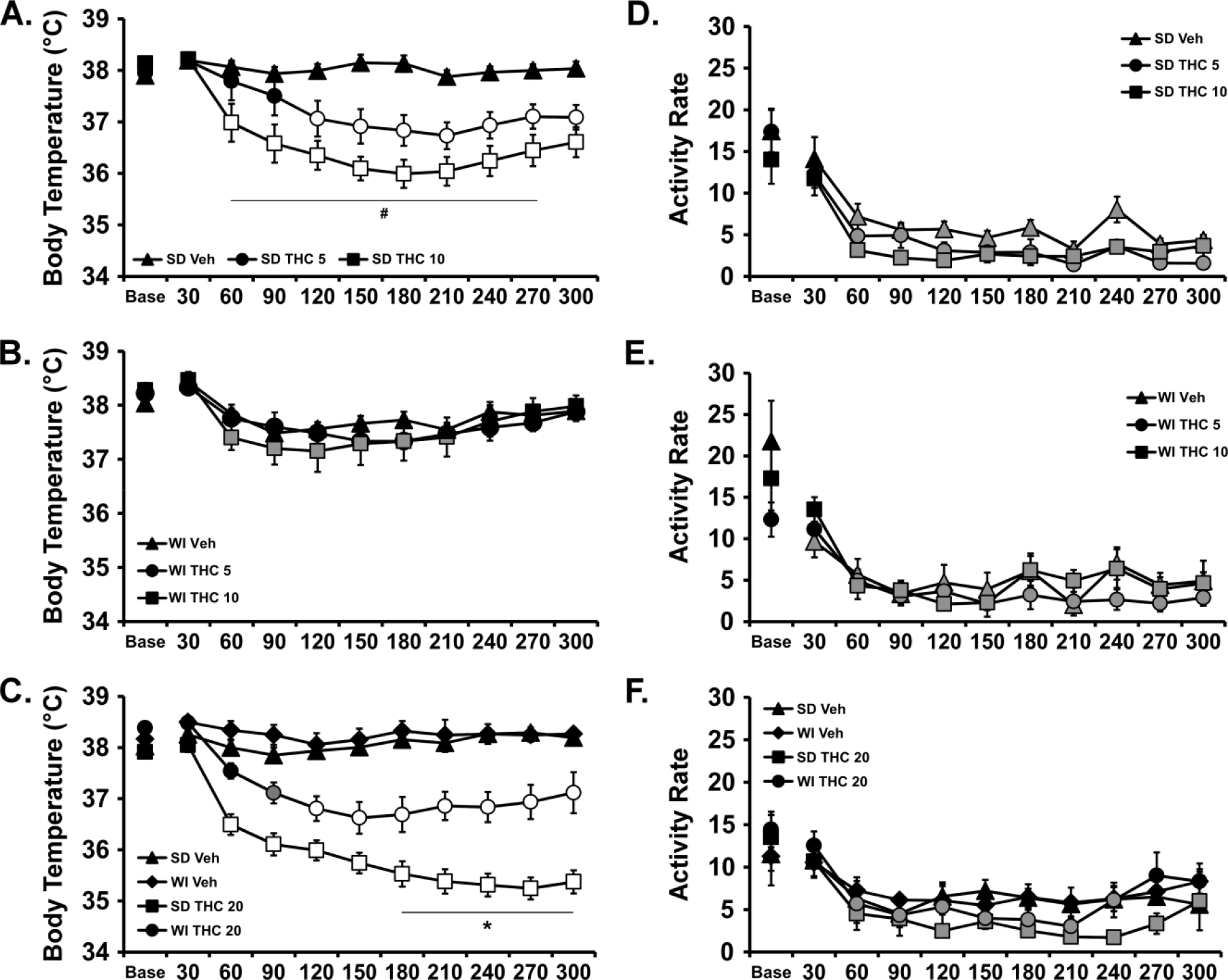
Mean (N=8 / group; SEM) body temperature (A, B, C) and activity rate (D, E, F) following i.p. injection with THC. A,D) Sprague-Dawley and B,E) Wistar groups after injection with THC (0.0, 5.0, 10.0 mg/kg) and C,F) both groups after injection with THC (0.0, 20.0 mg/kg). Base=baseline. Open symbols indicate a significant difference from both vehicle at a given time-point and the within-treatment baseline, while shaded symbols indicate a significant difference from the baseline only. A significant difference from the 5.0 dose at a given time is indicated by # and between strains at a given time for the same dose by *.

Analysis of the activity rates confirmed a significant effect of Time after injection [F (10, 420) = 73.87; P<0.0001] but not of Drug treatment / Group or any interaction of the factors (**Figure D,E**).

Since there was no effect of 10 mg/kg, THC, i.p., on temperature in the WI group, a followup study was conducted to evaluate the effect of 20 mg/kg THC, i.p. For this, rats within each group were injected with THC (0.0, 20 mg/kg, i.p.) in counter-balanced treatment order and significant THC-related hypothermia was produced in each strain (**Figure 4C**). The three-way ANOVA confirmed main effects of Drug Condition [F (1, 308) = 598.2; P<0.0001], of Strain [F (1, 308) = 105.7; P<0.0001], of Time Post-Injection [F (10, 308) = 16.71; P<0.0001] and of the interactions of Time with Drug Condition [F (10, 308) = 15.41; P<0.0001] and of Strain with Drug Condition [F (1, 308) = 59.35; P<0.0001], on the body temperature. The post-hoc test failed to confirm any differences in body temperature between strains, nor any differences relative to baseline within either strain, following vehicle injection. Body temperature was significantly lower than the baseline after 20 mg/kg THC, i.p., in both Wistar (90-300 minutes post-injection) and Sprague-Dawley (60-300 minutes post-injection) groups and significantly lower compared with respective timepoints after vehicle injection for both Wistar (120-270 minutes post-injection) and Sprague-Dawley (60 - 300 minutes post-injection) groups. Finally, the body temperature was lower in the Sprague-Dawley rats compared with the Wistar rats from 180 to 300 minutes post-injection.

The three-way ANOVA confirmed main effects of Drug Condition [F (1, 308) = 5.85; P<0.05], of Strain [F (1, 308) = 5.34; P<0.05] and of Time Post-Injection [F (10, 308) = 10.83; P<0.0001] on activity rate (**Figure 4D**). The post-hoc test failed to confirm any differences inactivity between strains within dosing condition, or differences attributable to THC within strain, at a given timepoint after injection.

### 3.4 Experiment 4 (SR141716 prior to vapor inhalation)

The SR (4 mg/kg, i.p.) failed to significantly alter the body temperature responses to inhaled THC in the second vapor study (**Figure 5A, C**). In the Wistar group, the ANOVA confirmed significant effects of Time after vapor initiation [F (8, 192) = 17.93; P<0.0001] and the interaction of Time with Drug Condition [F (24, 192) = 2.43; P<0.0005] on body temperature. The post-hoc test confirmed that body temperature was significantly reduced by THC inhalation relative to the respective pre-treatment/PG inhalation conditions and failed to confirm any difference between vehicle and SR pre-treatment conditions for either THC or PG inhalation conditions. The analysis for the Sprague-Dawley rats confirmed significant effects of Time [F (8, 192) = 24.17; P<0.0001], Drug Condition [F (3, 24) = 5.78; P<0.005] and the interaction of Time with Drug Condition [F (24, 192) = 4.61; P<0.0001] on body temperature. Again, the post-hoc test confirmed that body temperature was significantly reduced by THC inhalation relative to the respective pre-treatment/PG inhalation conditions and failed to confirm any difference between vehicle and SR pre-treatment conditions for either THC or PG inhalation conditions.

**Figure 5:**
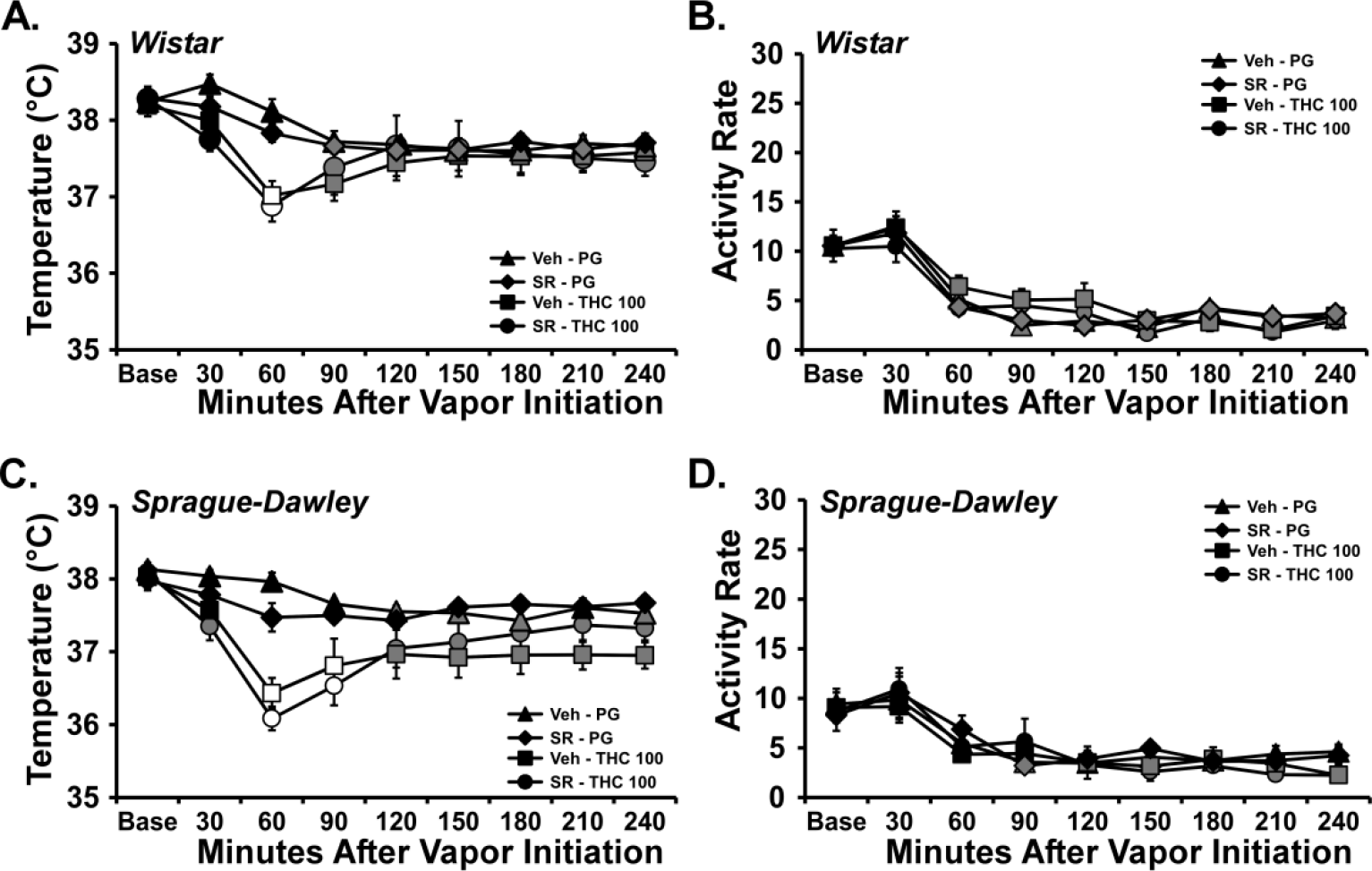
Mean (N=7 / group; SEM) body temperature following vapor inhalation of the PG vehicle or THC (100 mg/mL in the PG) for 30 minutes. Treatment conditions included an injection of vehicle or SR141716 (SR; 4 mg/kg, i.p.) 15 minutes prior to vapor initiation. Open symbols indicate a significant difference from both PG (within respective pre-treatment) at a given time-point and the within-treatment baseline, while shaded symbols indicate a significant difference from the baseline only. A difference from the PG condition (within respective pre-treatment, but not the baseline, is indicated by *. Base: Baseline 30 minutes prior to vapor initiation. SR: SR141716; Veh: Vehicle.

Activity was significantly affected by time after the start of inhalation but not by the dosing condition in WI [Time: F (8, 192) = 66.53; P<0.0001] and SD [Time: F (8, 224) = 20.17; P<0.0001] rat groups (**Figure 5 B, D**).

The vehicle pre-injection conditions were analyzed together to compare the strain response to THC as a replication of the first study. In this case the three-way ANOVA confirmed main effects of Inhalation Condition [F (1, 216) = 70.11; P<0.0001], of Strain [F (1, 216) = 31.03; P<0.0001], of Time after initiation of inhalation [F (8, 216) = 12.58; P<0.0001] and of the interactions of Time with Drug Condition [F (8, 216) = 4.224; P<0.0005] and of Strain with Drug Condition [F (1, 216) = 8.62; P<0.005], on the body temperature. The post-hoc test did not confirm any signficant differences between strains at any specific time after PG or THC inhalation, however.

### 3.5 Experiment 5 (Anti-nociception)

Anti-nociceptive effects of inhaled THC were produced in each strain of rats as illustrated in **Figure 6**. The ANOVA confirmed a significant effect of Drug inhalation condition [F (2, 28) = 14.45; P<0.0001], but not of Strain or of the interaction of factors, on tail withdrawal latency. The post-hoc test found that latency, collapsed across strain, was longer following THC (200 mg/mL) inhalation compared with either PG or THC (100 mg/mL) inhalation.

**Figure 6:**
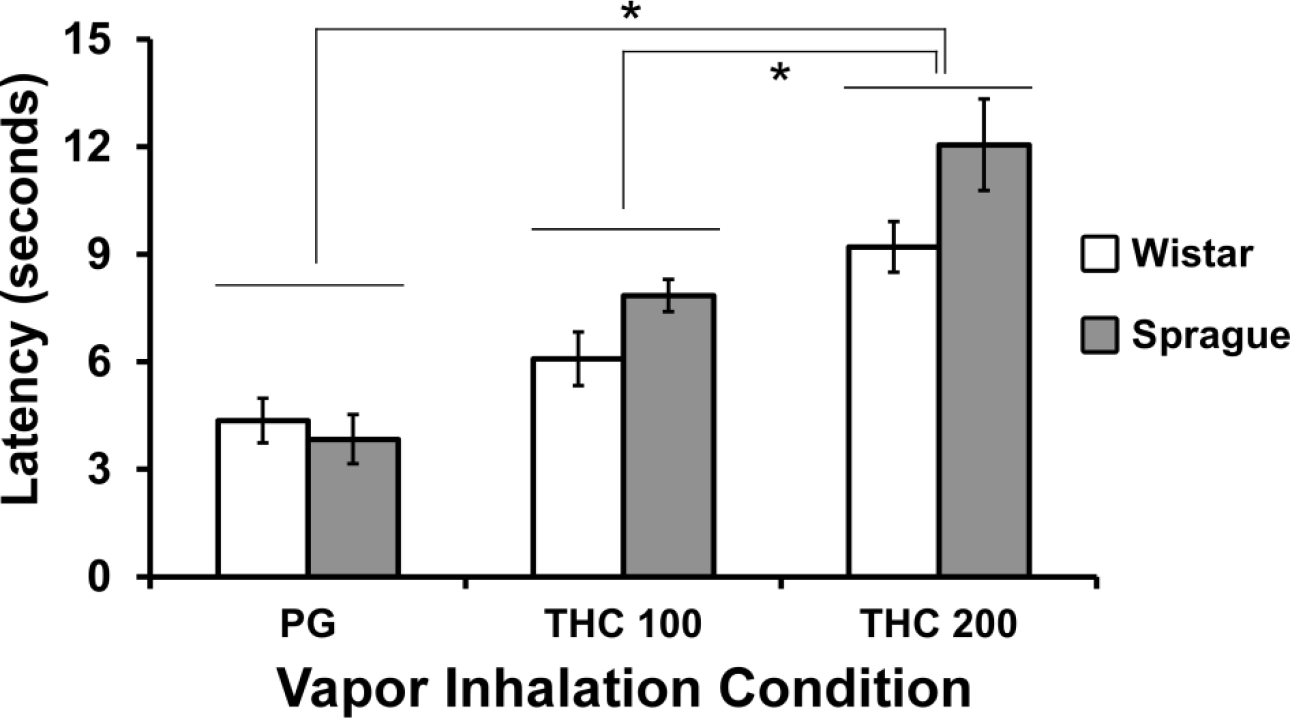
Mean (N= 8 / group; SEM) tail withdrawal latency following vapor inhalation of the PG vehicle or THC (100, 200 mg/mL in the PG) for 30 minutes. A significant difference between vapor conditions, collapsed across strain, is indicated by *.

### 3.6 Experiment 6 (Plasma THC 3,4)

The inhalation of THC (50, 200 mg/mL in the PG) for 30 minutes resulted in identical plasma THC between the two groups of rats (n.s. effect of Strain and the interaction of Strain with Dose condition) when assessed at 33-34 weeks of age (**Figure 7A**), but a significantly higher plasma THC concentration was observed with the 200 mg/mL vs 50 mg/mL concentration (F (1, 18) = 38.63; P<0.0001). Analysis of the plasma THC concentrations observed in the i.p. injection study failed to confirm any significant effects of strain, dose or the interaction of factors (**Figure 7B**).

**Figure 7:**
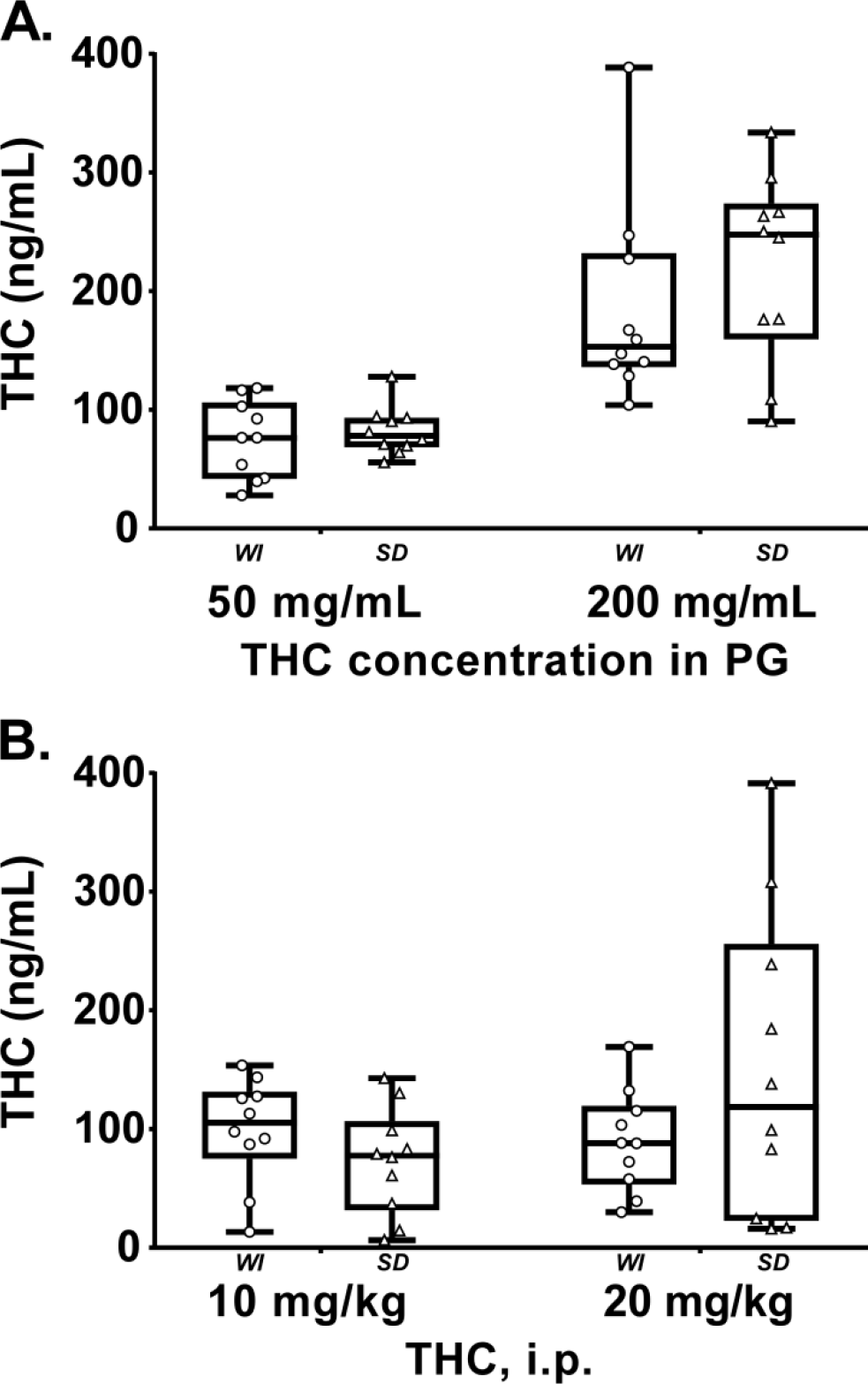
Plasma THC concentrations for Wistar (WI) and Sprague-Dawley (SD) rats (N=10 / group) following A) vapor inhalation of the THC (50, 200 mg/mL) for 30 minutes or B) THC injection (10, 20 mg/kg, i.p.). Box plots depict median and interquartile range and the bars indicate the range. Individual subject values are also plotted.

### 3.7 Experiment 7 (SR141716 prior to THC i.p.)

The 4 mg/kg SR dose significantly altered the body temperature responses to injected THC in each strain (**Figure 8**). In the Wistar group the 30 mg/kg, i.p. THC dose significantly reduced body temperature but this was prevented by SR injected 15 minutes prior to the THC [Drug condition: F (3, 21) = 5.23; P<0.01; Time post-injection: F (10, 70) = 4.94; P<0.0001; Interaction: F (30, 210) = 7.17; P<0.0001].

**Figure 8:**
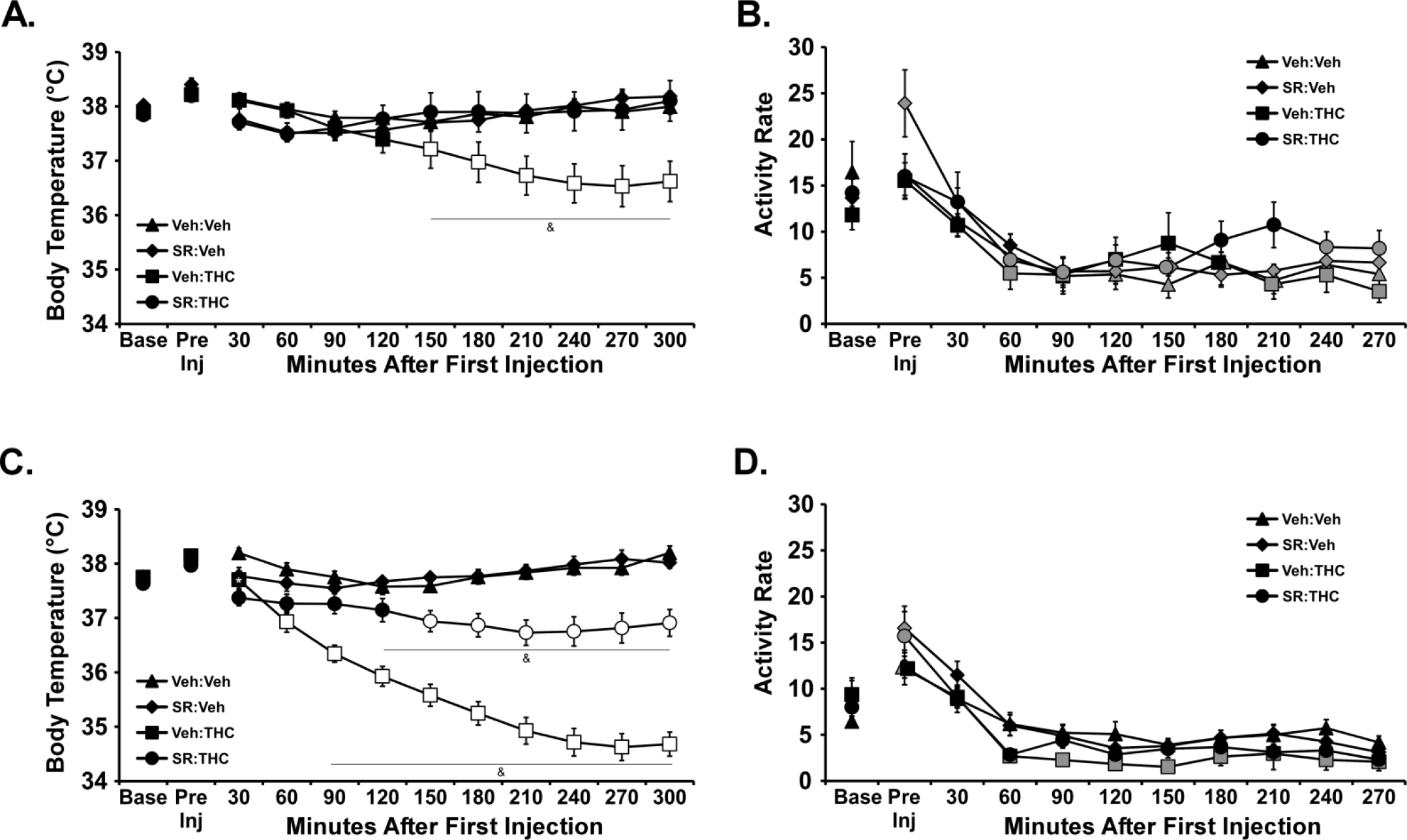
Mean (N= 6 / group; SEM) body temperature following injection of THC in A) Wistar (0, 30 mg/kg, i.p.) or B) Sprague-Dawley (0, 20 mg/kg, i.p.) rats. Treatment conditions included an injection of vehicle or SR141716 (SR; 4 mg/kg, i.p.) 15 minutes prior to the THC injection. Open symbols indicate a significant difference from both vehicle (within respective pre-treatment) at a given time-point and the within-treatment baseline, while shaded symbols indicate a significant difference from the baseline only. A significant difference from all other treatment conditions is indicated with &. A significant difference from the vehicle condition (within respective pre-treatment), only, is indicated by *. Base: Baseline 30 minutes prior to vapor initiation. SR: SR141716; Veh: Vehicle.

Similarly, in the Sprague-Dawley group the 20 mg/kg, i.p. THC dose significantly reduced body temperature but this was attenuated by SR injected 15 minutes prior to the THC [Drug condition: F (3, 21) = 56.72; P<0.0001; Time post-injection: F (10, 70) = 65.74; P<0.0001; Interaction: F (30, 210) = 29.92; P<0.0001].

Activity was significantly altered by Time Post-Injection [F (10, 280) = 30.84; P<0.0001] and the interaction of Time with Drug condition [F (30, 280) = 1.60; P<0.05] in the Wistar group but only by Time Post-Injection [F (10, 280) = 48.73; P<0.0001] in the Sprague-Dawley rats. The post-hoc tests confirmed that activity rate was higher in the SR-Vehicle condition compared with all other drug conditions after the pre-injection in Wistar rats (and compared with the SR-Vehicle pre-injection timepoint in Sprague-Dawley rats) but no other treatment-related differences between or within either group.

### 3.8 Experiment 8 (THC vapor inhalation for 40 minutes)

The inhalation of THC (100 mg/mL) for 40 minutes significantly reduced the body temperature of each rat strain at 55-56 weeks of age (**Figure 9**). Data collection for two rats of each strain was inadvertently terminated after the 150 minutes post-initiation. Initial analysis through this timepoint with the full sample revealed no differences in interpretation thus these individuals were omitted to preserve the repeated-measures analysis of the entire time-course. The three-way ANOVA confirmed main effects of Inhalation Condition [F (1, 160) = 128.2; P<0.0001], of Strain [F (1, 160) = 115.7; P<0.0001], of Time after initiation of inhalation [F (7, 160) = 8.52; P<0.0001] and of the interactions of Time with Inhalation Condition [F (7, 160) = 6.73; P<0.0001] and of Strain with Drug Condition [F (1, 160) = 52.55; P<0.0001], on the body temperature.

The activity rate was unchanged by THC inhalation although it was increased in the Wistar group by the onset of the vapor whether PG or THC. The three-way ANOVA confirmed main effects of Time [F (7, 160) = 19.18; P<0.0001], of Strain [F (1, 160) = 32.18; P<0.0001] and of the interactions of Time with Strain [F (7, 160) = 2.36; P<0.05]. The post hoc test further confirmed that activity rate was increased over the baseline rate for the WI animals 60 minutes after the start of inhalation and also increased relative to the SD rats at the same time after inhalation of either THC or PG, respectively.

**Figure 9:**
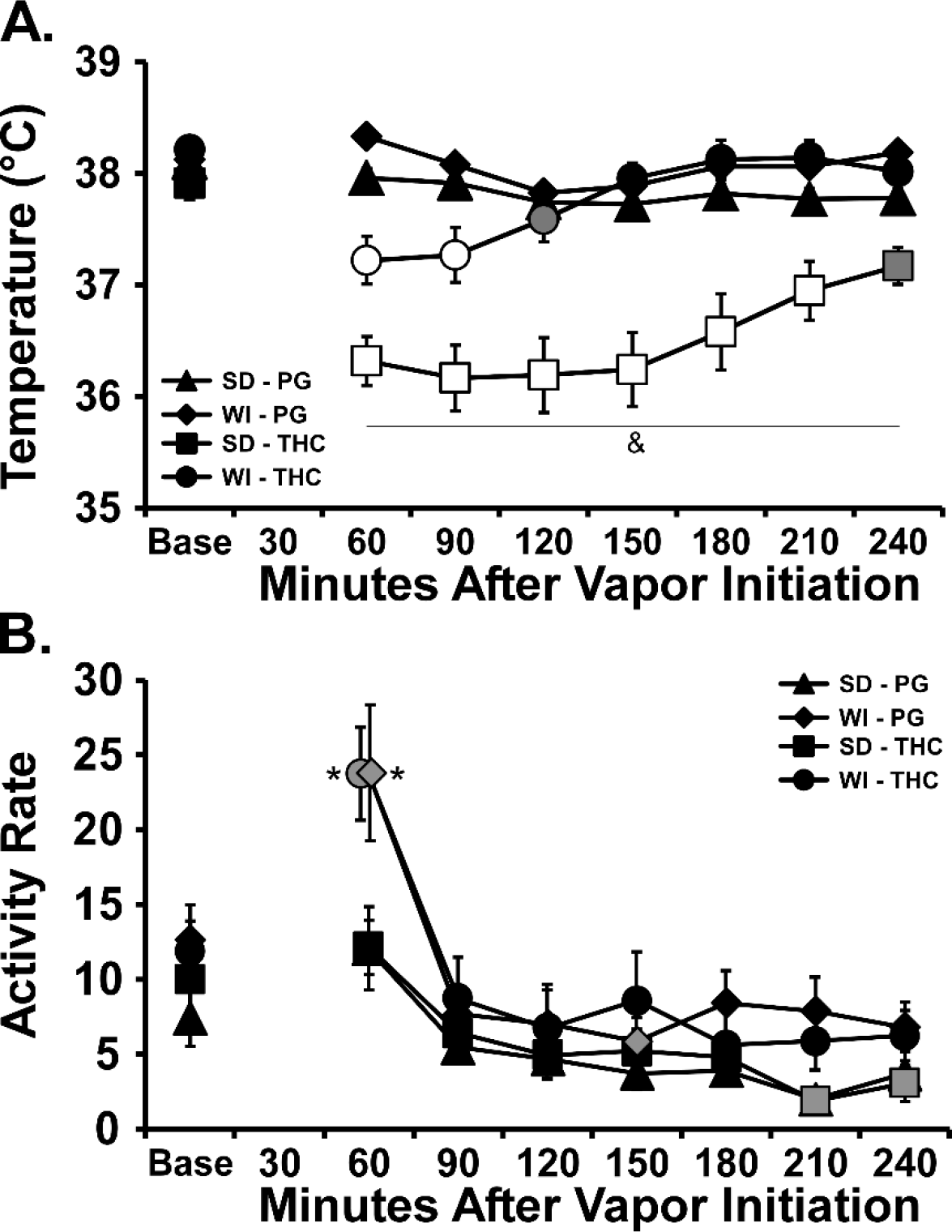
Mean (N= 6 / group; SEM) A) body temperature and B) activity rate after vapor inhalation of THC (100 mg/mL) for 40 minutes. Open symbols indicate a significant difference from both PG (within respective pre-treatment) at a given time-point and the within-treatment baseline, while shaded symbols indicate a significant difference from the baseline only. A significant difference from all other treatment conditions at a given time point is indicated with & and a difference between strains for a given time and inhalation condition with *.

## 4. Discussion

This study shows that the inhalation of THC vapor using an e-cigarette approach induces hypothermia in both Wistar and Sprague-Dawley male rats, with the latter strain exhibiting a significantly greater reduction in temperature after identical inhalation conditions. This work was motivated by an indirect observation of a hypothermic insensitivity of Wistar rats relative to Sprague-Dawley rats in our prior vapor inhalation studies (Javadi-Paydar et al. 2018a; Nguyen et al. 2016) as well as a similar outcome of preliminary pilot studies for an investigation of the impact of CBD co-administration with THC (Taffe et al. 2015). The difference in the present study was observed in age-matched, identically treated groups and the difference was present in the initial THC inhalation exposure, in the intraperitoneal injection studies and in the final vapor inhalation investigation across an ~one year interval. The data therefore confirm a quantitative, but not qualitative, strain difference in the thermoregulatory response to inhaled (and injected) THC without a difference in the anti-nociceptive effect. Since the Sprague-Dawley rats were significantly smaller in size, it was critical to determine if THC dose differed between strains under the inhalation conditions. The pharmacokinetic data show that identical plasma concentrations of THC were produced in each group after 30 minutes of inhalation of THC vapor at the 100 mg/mL concentration. Thus, the strain difference in body temperature response cannot be attributed to the delivery of a different dose and strain differences must therefore be attributable to other factors.

The consistency of THC plasma levels following vapor inhalation across rats that differed in body size, due to strain, echoes the result of a prior study which showed approximately equivalent plasma concentrations of THC after inhalation in male and female Wistar rats that differed significantly in body weight by sex (Javadi-Paydar et al. 2018a). In that prior study, however, male and female Wistar rats exhibited a similar magnitude of hypothermia and anti-nociception produced by the THC. The difference in plasma THC across the two observation timepoints in this study was unexpected, particularly given the similarity of the temperature response across the first and second vapor inhalation experiments (Figure 2, 5). However, this is the first time we have explicitly compared plasma levels of THC across significant intervals of time in the same rats following inhalation. Additional experimentation would be required to determine if this is due to different respiration / weight, different drug distribution, different metabolism or possibly differential behavior in the vapor exposure chamber. With respect to this latter, animals could vary where they put their nose, relative to the vapor delivery location and thereby change the exposure; inspection of the individual distributions in Figure 3 supports this possibility. Also of interest is that plasma THC concentrations reached following inhalation of THC at the 200 mg/mL concentration were not substantially higher compared with plasma concentrations after the 100 mg/mL dose which may reflect a ceiling on the rate of intrapulmonary uptake as drug concentration is increased. For the present purposes, however, it is most critical that there was no significant difference across the strains at any time of plasma collection.

There was no strain difference in the anti-nociception assay, which shows that the influence of strain on hypothermia is dissociable from the effects on anti-nociception. This dissociation combines with the similar plasma concentrations across strain to further confirm that the strain-related hypothermia difference was not due to different THC dose. It suggests instead that Wistar rats are more resistant to body temperature dysregulation than are Sprague-Dawley rats, either in response to THC or more generally. Consistent with this interpretation, one prior report shows a differential hypothermia in response to challenge with the serotonin receptor 1a agonist 8-OH-DPAT in different breeding colony sources of Sprague-Dawley rats and in Wistar rats (Kelder and Ross 2001). The data also appeared to suggest a more pronounced strain difference in thermoregulatory sensitivity after i.p. injection compared with the difference after inhalation of THC. This may possibly reflect differences in THC distribution throughout the larger body mass of the Wistar rats, in part or in whole, indeed strain differences in THC-induced hypothermia appeared to grow more pronounced as the animals aged and the body size difference increased. This conclusion seems unlikely, however, given that the strains did not differ in plasma concentrations of THC when exposed to identical inhalation conditions or when administered an equivalent mg/kg dose by i.p. injection. One alternate hypothesis is that Wistar rats became more tolerant from a pharmacodynamic standpoint, compared with the Sprague-Dawley rats, in the course of the repeated (albeit intermittent) exposure. Recent work with the inhalation model shows that adult WI male rats do not become tolerant to THC after 4 sequential days of twice-daily vapor inhalation (Nguyen et al. 2018), thus it seems unlikely that there would be significant pharmacodynamic tolerance with the present dosing history. Together the evidence suggests the apparent strain difference in thermoregulatory response to THC that interacted with route of administration is not directly related to THC exposure but may be related to non-cannabinoid mechanisms involved in temperature regulation.

There was no evidence of any strain differences in locomotor suppression caused by THC in this study but conclusions must be tempered by the general failure to observe THC-related effects on activity. This is somewhat expected since activity rate assessed by radiotelemetry was only inconsistently changed by THC inhalation in our prior studies using these methods (Javadi-Paydar et al. 2018a; Nguyen et al. 2016) and it required a 30 mg/kg, i.p. injection dose in male Sprague-Dawley rats to suppress locomotion consistently in another study (Taffe et al. 2015).

One curious finding in this study was that a dose of the CB1 antagonist/inverse agonist SR141716 which was sufficient to block (WI) or attenuate (SD) the body temperature response to THC administered intraperitoneally (also see Taffe et al. 2015) did not significantly alter the response to vapor inhalation of THC. SR141716 did not appear to alter the initial nadir in body temperature observed 60 min after the start of inhalation but may perhaps have slightly facilitated the return to normative temperature after about 90 (WI) or 120 (SD) minutes after the initiation of vapor inhalation. Interestingly the body temperature of rats only started to diverge from that observed with vehicle pre-injection 60 (SD) or 150 (WI) minutes after THC i.p. injection. This finding warrants additional followup study, particularly since it is consistent with preliminary results we have reported in a pre-print for female Wistar rats (Nguyen et al. 2017).

Overall this study further confirms the efficacy of a new electronic-cigarette based method of delivering THC to rats and shows the effects are qualitativedly similar in age-matched groups of two widely used strains of laboratory rat. Importantly, Wistar rats appear less prone to hypothermia compared with Sprague-Dawley males when administered an equivalent THC dose. The potential for these quantitative differences should be taken into account when attempting to replicate previously published methods in different rat strains.

## Author Contributions

MAT designed the studies, with refinements contributed by KMC, SAV and TMK. MC designed the vapor inhalation equipment that was used. KMC, SAV and TMK performed the research and conducted initial data analysis. MAT conducted statistical analysis of data, created figures and wrote the initial draft of the paper. All authors approved of the submitted version of the manuscript.

## Acknowledgements

The authors are greatful to Shawn Aarde, Ph.D. for contributions to the invention and validation of this method of drug delivery for rats and to Mehrak Javadi-Paydar, Ph.D. and Jacques Nguyen, Ph.D. for ongoing discussion and input on the use of the model. This is manuscript #29709 from The Scripps Research Institute.

## Abbreviations

PG: propylene glycol
THC: Δ^9^tetrahydrocannabinol

